# Towards establishing no observed adverse effect levels (NOAEL) for different sources of dietary phosphorus in feline adult diets: results from a 7 month feeding study

**DOI:** 10.1101/2020.09.01.276907

**Authors:** Jennifer C. Coltherd, Janet Alexander, Claire Pink, John Rawlings, Jonathan Elliott, Richard Haydock, Laura J. Carvell-Miller, Vincent Biourge, Luis Molina, Richard Butterwick, Darren W. Logan, Phillip Watson, Anne Marie Bakke

**Affiliations:** Waltham Petcare Science Institute, Melton Mowbray, Leicestershire, LE14 4RT, UK; Department of Comparative Biomedical Sciences, Royal Veterinary College, University of London, London NW1 0TU, UK; Royal Canin SAS, 650 Avenue de la Petite Camargue, 30470 Aimargues, France

## Abstract

High dietary phosphorus (P), particularly soluble salts, may contribute to chronic kidney disease development in cats. The aim of this study was to assess the safety of P supplied at 1g/1000kcal from a highly soluble P salt in P-rich dry format feline diets. Seventy-five healthy adult cats (n=25/group) were fed either a low P control (1.4g/1000 kcal; calcium:phosphorus ratio, Ca:P 0.97) or one of two test diets with 4g/1000 kcal (4184kJ); Ca:P 1.04 or 5g/1000kcal; Ca:P 1.27, both incorporating 1g/1000kcal (4184kJ) sodium tripolyphosphate (STPP) – for a period of 30 weeks in a randomised parallel-group study. Health markers in blood and urine, glomerular filtration rate, renal ultrasound and bone density were assessed at baseline and at regular time points. At the end of the test period, responses following transition to a commercial diet (total P – 2.34g/1000kcal, Ca:P 1.3) for a 4-week washout period were also assessed. No adverse effects on general, kidney or bone (skeletal) function and health were observed. P and Ca balance, some serum biochemistry parameters and regulatory hormones were increased in cats fed test diets from week 2 onwards (p≤0.05). Data from the washout period suggest that increased serum creatinine and urea values observed in the two test diet groups were influenced by dietary differences during the test period, and not indicative of changes in renal function. The present data suggest NOAELs for feline diets containing 1g P/1000kcal (4184kJ) from STPP and total phosphorus level of up to 5g/1000kcal (4184kJ) when fed for 30 weeks.

## Introduction

Chronic kidney disease (CKD) has long been established as the most prevalent metabolic disease and the leading cause of death in domestic cats over the age of 12 years^(1; 2)^. It is a progressive disease that can be triggered by a number of genetic and environmental factors^(3)^. Among acquired causes of the disease, dietary phosphorus (P) has been linked to its development and/or progression, not only in cats^(2)^, but also in humans^(Reviewed in 4)^.

Several studies have indicated that inclusion of high levels of soluble P salts in diets, often added for technical, palatability and nutritional reasons, can cause glucosuria and increase blood urea nitrogen content whilst decreasing creatinine clearance^(5; 6)^. In a recent feeding study^(2)^, reduced feed intake and vomiting were observed within four weeks in adult cats fed high dietary levels of sodium dihydrogen phosphate (NaH_2_PO_4_; SDHP) providing 3.6g/1000kcal (4184kJ) P (total P 4.8g/1000kcal; calcium:phosphorus ratio [Ca:P] 0.6). This led to structural and functional changes in the kidneys as indicated by reduced GFR and proteinuria, compatible with early stage CKD.^(2)^ This may have been due to the high post-prandial serum phosphate levels induced by the test diet^(7)^. In a subsequent 28-week study, exposure to a more moderate level of SDHP, providing 1.5g/1000kcal P (total P 3.6g/1000kcal; Ca:P 0.9) resulted in the development of renoliths and/or structural changes in the kidneys^(2)^. Although biochemical markers of kidney health remained within physiological reference ranges in most of these cats, three out of 25 in the test group were removed from the study due to persistent azotaemia^(2)^.

A no observed adverse effect level (NOAEL) has therefore yet to be established for total P or the inclusion of soluble P salts, and currently no specific nutritional guidance for safe upper limits for dietary P exist for feline diets. Since 2018, however, The European Pet Food Industry FEDIAF, has added a precautionary footnote to maximum dietary P recommendations, suggesting that inorganic P compounds may pose a risk to feline renal function^(2; 8; 9)^. The studies reported by Alexander et al.^(2)^ indicate that further long term feeding trials are needed to determine safe upper limits of dietary P and particularly in the form of added soluble salts. Dietary P from organic raw materials, such as meat and bone meals or plant-based ingredients, has been shown to be less bioavailable than from soluble P salts^(6; 10)^. Recent studies in healthy adult cats investigating post-prandial responses following single meal exposure indicate that soluble P salts, but not organic P sources, induce rapid, dose-dependent increases in serum phosphate and parathyroid hormone (PTH), a key regulator of mineral homeostasis^(7)^. There is also evidence that increasing the Ca:P may mitigate the effects of exposure to high levels of soluble P salts in the diet^(7; 11)^. Current FEDIAF guidance for feline diets recommends Ca:P between 1:1 and 2:1^(12)^.

To provide further evidence to inform a safe upper limit for soluble P inclusion in cat diets, a 30 week feeding study was initiated to investigate the effects of 1g P/1000kcal (4184kJ) of sodium tripolyphosphate (Na_5_P_3_O_10_, STPP). Two diets were tested at this level, containing either 4g/1000kcal (4184kJ) or 5g/1000kcal (4184kJ) levels of total P, with Ca:P ratios of 1 and 1.3, respectively. This enabled two hypotheses to be tested, firstly that a diet formulated to 4g/1000kcal (4184kJ) total P (incorporating 1g P from STPP) and a Ca:P =1 would induce no observable adverse effects; and secondly increasing the total P to 5g/1000kcal (4184kJ), maintaining the STPP contribution at 1g, and increasing Ca:P to 1.3 would also induce no observable effects and allow more flexibility in dietary formulations using organic raw materials. To evaluate safety, markers of renal and bone function, as well as Ca and P homeostasis were assessed throughout the course of the study.

## Materials and Methods

This work was approved by the WALTHAM Animal Welfare and Ethical Review Body and the local Institutional Animal Care and Use Committee (IACUC, #ONL 19-001).

### Animals and husbandry

Adult cats housed at an independent research facility were health-screened based on assessment of serum biochemistry, haematology and urinary health parameters, as well as Dual-energy X-ray absorptiometry (DXA) and abdominal ultrasound scans. Prior to screening, the cats had been fed one of two commercial feline diets, one wet and one dry format (see table 1 for analysis), for varying amounts of time (1-4 months). Those with findings outside of the respective laboratory reference intervals or otherwise exhibiting signs of ill health as considered by the site veterinarian were excluded, as were cats identified as having pre-existing uroliths, renoliths or abnormal kidney architecture (i.e. extremely small kidneys or high echogenicity) as determined by an independent specialist in veterinary diagnostic imaging.

**Table 1.** Experimental diet composition in g/1000kcal (4184kJ) or g/100g.

| Nutrient | Control |  | Moderate |  | High |  | Screening diet: Wet<sup>‡</sup> |  | Screening and Washout diet: Dry<sup>§</sup> |  |
| --- | --- | --- | --- | --- | --- | --- | --- | --- | --- | --- |
|  | (g/1000kcal) | (g/100g) | (g/1000kcal) | (g/100g) | (g/1000kcal) | (g/100g) | (g/1000kcal) | (g/100g) | (g/1000kcal) | (g/100g) |
| Phosphorus | 1.43 | 0.55 | 4.03 | 1.49 | 5.03 | 1.84 | 3.61 | 0.36 | 2.34 | 0.86 |
| Inorganic P present (source) | 0.06 | 0.02 | 1.0 | 0.37 | 1.0 | 0.38 | 0.9 | 0.09 | 0.78 | 0.3 |
|  | (palatant) |  | (STPP) |  | (STPP) |  | (STPP) |  | (palatant) |  |
| Calcium | 1.38 | 0.53 | 4.20 | 1.55 | 6.38 | 2.34 | 3.13 | 0.31 | 3.03 | 1.11 |
| Ca:P | 0.96 | 0.96 | 1.04 | 1.04 | 1.27 | 1.27 | 0.90 | 0.90 | 1.30 | 1.30 |
| Sodium | 2.00 | 0.77 | 2.01 | 0.74 | 2.12 | 0.78 | 2.73 | 0.27 | 1.27 | 0.47 |
| Magnesium | 0.23 | 0.09 | 0.26 | 0.10 | 0.31 | 0.11 | 0.19 | 0.02 | 0.23 | 0.09 |
| Chloride | 3.96 | 1.52 | 1.76 | 0.65 | 1.69 | 0.62 | 1.30 | 0.13 | 2.01 | 0.74 |
| Vitamin D<sub>3</sub><sup>*</sup> | 484.00 | 186.00 | 436.23 | 161.00 | 493.43 | 181.00 | 221.26 | 22.10 | 171.20 | 63.00 |
| Protein | 80.15 | 30.80 | 90.36 | 33.35 | 98.55 | 36.15 | 97.01 | 9.69 | 78.64 | 28.94 |
| Fat | 35.65 | 13.70 | 34.14 | 12.60 | 35.30 | 12.95 | 75.09 | 7.50 | 34.84 | 12.82 |
| Ash | 14.83 | 5.70 | 23.71 | 8.75 | 28.62 | 10.50 | 21.73 | 2.17 | 22.55 | 8.30 |
| Crude Fibre | 10.28 | 3.95 | 9.89 | 3.65 | 10.09 | 3.70 | 2.00 | 0.20 | 10.33 | 3.80 |
| Moisture | 15.22 | 5.85 | 17.77 | 6.56 | 17.26 | 6.33 | 812.94 | 81.20 | 18.13 | 6.67 |
| Predicted Metabolisable Energy, PME<sup>†</sup> |  | 384.30 |  | 369.1 |  | 366.8 |  | 99.9 |  | 368.0 |

<sup>\*</sup>International units IU/1000kcal or IU/100g <sup>†</sup>Calculated by proximate analysis to PME kcal/100g according to LaFlamme<sup>(15)</sup> <sup>‡</sup>Whiskas pate with Chicken <sup>§</sup>Royal Canin Indoor Dry (cat)

The seventy five cats included in the study (thirty eight neutered males, twenty seven neutered females and ten entire females) were aged between 1.9 and 8.6 years (median age 5.3 years) at the start of the study. Before initiation of the feeding study, cats were subjected to a stratified randomisation across diet groups to balance gender and neuter status, age, energy intake and body condition score (BCS). All cats were group housed within the colony across six different social rooms, capacity varied with room size and there were 20 cats in the largest group. Deionised water was offered *ad libitum* and social rooms are enriched with multi-level furniture, shelving and windows so that cats have access to more of the room and are able to view the surroundings. To enable individual feeding, cats received two meals per day, each providing 50% of their maintenance energy requirements (MER), for 30 minutes within their assigned lodges (measuring 122cm x 90cm x 76cm [height x length x depth], multileveled with hammocks and privacy barriers), arrangements that they were habituated to. When individual housing was required during urine and faeces collection phases, they were placed within the same assigned lodges, within their normal rooms, overnight. During the day, cats were group housed and monitored to enable socialization and normal behaviours during the collection periods.

Throughout the study period, diets were offered to maintain body weight within 10% of their initial weight and BCS assessed according to a 9-point scale^(13)^. Deionised water was offered *ad libitum* throughout the study.

### Study Design

The study assessed kidney health effects in adult cats offered one of three experimental diets during a 30-week feeding study in a parallel design followed by a 4-week washout period. The number of cats included was based on power analyses (detailed in Statistical Powering section). All cats were initially offered the control diet (Table 1) for 5 weeks. In the final 2 weeks of this pre-feed period, baseline blood, faeces and urine samples were collected. Following this, each group was then offered one of three experimental diets (Table 1) differing in P content, source and Ca:P ratios for 30 weeks. All cats were then transitioned onto the dry format commercial diet, fed during the screening phase, for the 4-week washout period, after which an additional blood and urine sample was collected (see Table 1 for diet analyses).

### Diets

Single batches of all three experimental dry format extruded diets were manufactured by Mars Petcare North America (Franklin, Tennessee, USA) using ingredients routinely used in commercial cat foods. All diets were produced through a standard extrusion process common in the pet food industry^(14)^. For ease, these experimental diets will subsequently be referred to as control, moderate and high throughout (see table 1 for analyses). The control diet, was formulated to meet the Association of American Feed Control Officials (AAFCO) and FEDIAF minimum recommended total P level of 0.5g/100 g dry matter (or 1.25g/1000kcal [4184kJ] based on an estimated MER of 100 kcal/kg^0.67^ body weight for adult cats) and a Ca:P ratio of 1.0. The P content of this diet was almost entirely supplied by the animal and plant derived raw materials, although a small contribution (0.06g P/1000kcal representing 4.4% of the total P) came from a commercial palatant. The two test diets both included sodium tripolyphosphate (Na5P3O10; STPP), at a level to provide a P at 1.0g/1000kcal (4184kJ). The test diets differed, however, in total P and Ca:P ratios with the moderate test diet formulated to contain 1.5g/100 g DM (4g/1000kcal or 4184kJ) and a Ca:P ratio of 1.0, and the high test diet formulated to contain 1.8g/100 g DM (5g/1000kcal or 4184kJ) and a Ca:P ratio of 1.3 (table 1). Due to the Ca and P content of chicken and pork meals used in the diets, the high test diet’s Ca:P could not be adjusted to <1.3.

### Analyses

Diets were nutritionally analysed to ensure they were complete and balanced according to AAFCO guidelines. All nutritional chemical analyses were carried out using Association of Official Agricultural Chemists (AOAC) procedures at Mars Petcare North America Regional Laboratory (Franklin, Tennessee).

Intake was recorded on an individual basis as mass (g) of diet offered minus mass (g) of diet refused. Body weight and body condition score were recorded weekly using the same calibrated scale.

### Mineral Balance

#### Faecal mineral content and apparent digestibility

At baseline and after 27 weeks of feeding, five day total urine and faeces collections were carried out to determine macronutrient, Ca and P apparent digestibility and balance. Faeces were stored at −20°C in a sealed container until the 5-day pooled sample was collected and analysed at the Royal Canin Laboratory, Guelph, Canada. Faeces were dried, homogenised, the moisture crude fibre, ash, fat and protein content measured according to established AOAC methods. Inductively Coupled Plasma (ICP) atomic emission spectroscopy (AES) analysis of P and atomic absorption spectrometry (AAS) analysis of Ca were also performed on faecal samples. Apparent digestibility (%) was calculated to represent the fraction of the nutrients that were retained from the diet rather than being excreted in the faeces using the following formula:

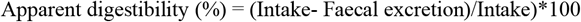

An estimation of the possible exposure of the cats to P and Ca was carried out using the formula:

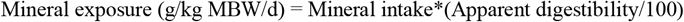

#### Urine mineral content and relative super saturation (RSS)

All urine excreted during a 5 day period was collected and stored at −80°C until analysis was performed. Urinary pH was assessed on freshly voided samples twice daily. The method for urine preparation and RSS analysis has been previously described^(16)^. Samples were analysed, at the Royal Canin Laboratory, Guelph, Canada, for oxalate, citrate, urate, potassium, calcium, sodium, ammonium, creatinine, sulphate, and phosphate via ion chromatography. The concentrations of minerals were then analysed by SUPERSAT software^(16)^ to calculate the RSS (activity product / solubility product) for struvite (magnesium ammonium phosphate; MAP) and Ca oxalate (CaOx). The urine concentration of P and Ca used for calculation of the urinary excretion of these minerals were determined via Inductively Coupled Plasma (ICP) analysis at the University of Guelph, Ontario, Canada.

Mineral balance (g) was also calculated for P and Ca as:

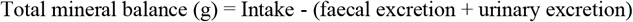

#### Urinalysis

At screening, baseline, 2, 4, 8, 12, 20, 27 weeks and following the 4 week washout period, a 5ml freshly voided sample was obtained from each cat for albumin, creatinine, protein, pH, and urine specific gravity analysis. Chemstrip 10A (Roche diagnostics) strips were employed with visual inspection via microscopy for urinary crystals and specific gravity was determined using a refractometer (ATAGO URC-Ne) within 30 minutes of collection. Following storage at −20°C, urine albumin and creatinine were measured using the Beckman Coulter Microalbumin (OSR6167) and Creatinine (OSR6178) assays for the Olympus AU480 biochemistry analyser (Olympus Europe GmbH). The urine albumin to creatinine ratio (UACR) was then calculated for each sample.

#### Blood based measures

At screening, baseline and week 2, 4, 8, 12, 20 and 28 of the study and following the 4 week washout period, morning fasted (>12 hrs) blood samples were collected for the measurement of standard biochemistry and haematology parameters. Vitamin D metabolites, parathyroid hormone (PTH), fibroblast growth factor 23 (FGF23), and markers of bone turnover – ionised calcium (iCa), serum cross laps (CTx) and bone specific alkaline phosphatase (BAP) were measured during the study and washout period only. Ethylenediaminetetraacetic acid (EDTA) anticoagulated blood was sent to IDEXX laboratories (Markham, Ontario Canada) for measurement of standard haematology parameters: white and red blood cell counts (WBC, RBC), haemoglobin concentration, haematocrit, platelet count, mean corpuscular volume (MCV), mean corpuscular haemoglobin (MCH), number and % of lymphocytes, monocytes and granulocytes. Serum was also prepared using serum separator tubes, these were stood upright at room temperature for a minimum of 30 minutes prior to a 10 minute centrifuge at 2000g, and sent to IDEXX laboratories (Markham, Ontario Canada) for determination of standard biochemistry parameters. These included total protein, albumin, phosphate, alkaline phosphatase (ALP), alanine transaminase (ALT), aspartate aminotransferase (AST), calcium, cholesterol, blood urea nitrogen (BUN), creatinine, triglycerides, and glucose, as well as symmetric dimethylarginine (SDMA, was not measured during screening). Further serum was prepared in the same manner for the measurement of BAP and CTx using the BAP MicroVue™ Quidel ELISA (TECO medical Group) and CartiLaps^®^ ELISA (Immunodiagnostic Systems Ltd), according to the manufacturer’s instructions; both assays have been previously validated for use in feline samples^(17)^. EDTA anticoagulated blood was centrifuged at 2000g for 10 minutes and plasma aliquoted, this plasma was sent to the Department of Comparative and Biomedical Sciences, Royal Veterinary College (London) for measurements of intact plasma FGF-23 and PTH concentration. FGF-23 was analysed using a sandwich ELISA (Kainos Laboratories Inc., Japan) as detailed by Geddes *et al*. 2013^(18)^, and PTH concentrations by a total intact PTH immunoradiometric assay (Scantibodies Laboratory, Inc. CA USA), previously validated for use with feline samples^(19)^. Serum was also prepared as above for the measurement of the vitamin D metabolites at the Bioanalytical Facility at the University of East Anglia, Norwich, UK using methods previously validated for cats^(2)^. Total 25OHD and total 24,25 (OH)_2_D were measured by LC/MS-MS (performed using a Micromass Quattro Ultima Pt mass spectrometer (Waters Corp., Milford, MA). Total serum 1,25 (OH)_2_D levels were measured using an EIA Kit (IDS Ltd., Tyne & Wear, UK). Ionised calcium, iCa, analysis was conducted using heparinised whole blood in a Stat Profile Prime Critical Care analyser (Nova Biomedical, MA, USA).

Serum calcium-phosphorus (CaP) product (mmol^2^/L^2^) was calculated by multiplying plasma total calcium and inorganic phosphorus concentrations in mmol/L.

#### Renal functional measurements

At baseline and week 4, 12, 20 and 28 of the study, iohexol clearance tests were carried out to enable an estimate of glomerular filtration rate using the method described by Finch *et al*.^(20)^. Briefly, 647 mg/kg iohexol (Omnipaque 300, Amersham Health, NJ, USA) was administered over a 2-3 minute period, via a cephalic catheter, followed by a heparinised saline flush (100 IU/ml, Wockhardt UK Ltd. Wrexham, U.K). The completion of the infusion represented time zero. Blood samples (1 mL) were collected from the cephalic vein into serum separator tubes at 2, 3 and 4 h post infusion. Serum iohexol concentration at each time point was analysed using High Performance Capillary Electrophoresis (deltaDOT Ltd., London BioScience Innovation Centre, London). Weight adjusted clearance (mL/kg/min) was calculated by the slope of the concentration gradient over the three time points.

The fractional excretion of P (FE_P_) and Ca (FE_Ca_) were calculated as the percentage filtered by the glomerulus and excreted into the urine, expressed as a ratio to creatinine clearance as below:

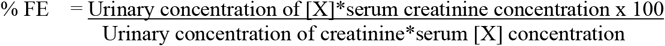

#### Imaging

At screening and at the end (week 30) of the study, a general physical health examination, DXA and abdominal ultrasound scans were carried out to detect soft tissue mineralization, urolithiasis or other pathology, with an additional ultrasound also performed during weeks 13 or 14. With the cats under sedation (6 mg/kg Intra Venous, IV, Propofol [10mg/mL]), radiographs of the abdomen were taken in ventrodorsal and right lateral recumbency, those of the thorax in right lateral recumbency and full abdominal ultrasound scans also carried out (6mHz probe on a Biosound Esaote MYLAB 30 machine). To confirm ultrasound findings at the end of the study (week 30), an additional enhanced resolution scan was performed with a 10-12mHz (linear probe) on a Logiq-E R7 machine with cats anaesthetised (0.14mg/kg intramuscular (IM) or subcutaneous (SQ) premedication premix consisting of Butorphanol (10mg/mL), Atropine (0.5mg/mL) and Acepromazine (10mg/mL), and anaesthetized with 5mg/kg IV Propofol (10mg/mL). All scans and radiograph interpretations were carried out by two independent specialist veterinary diagnostic imagers who were blinded to the dietary groupings.

#### Statistical powering

The study was powered according to the primary response variable GFR (iohexol clearance ml/min per kg), by simulation using baseline values from previous studies to estimate variance^(2)^. To detect a change in distribution where 10% of cats had values <0.92ml/min per kg, with approximately 80% power, 20 cats were required per diet group and an additional 5 cats per group were used in the study to allow for potential drop out.

#### Statistical analysis

Linear mixed effects models were used with log10 transformation where necessary and fixed effects of diet group, time point (weeks) and the interaction between diet group and time point. Individual cat was included as a random effect and time point was included as a factor, not a continuous variable. From these models mean values and 95% confidence intervals were estimated for each diet group and time point. For the pH measure, the median collection time was used for all estimates.

For the measures that had data for both screening and washout periods, a linear mixed effects model was fitted with the measure as the response, sampling time, diet group and their interaction as fixed effects and individual cat as a random effect. The estimates for each diet group at each time point were obtained from the model. The difference between washout and screening periods was calculated within each diet group using a Tukey honestly significantly different (hsd) test. These differences for each diet group were compared back to the control diet, again using a Tukey hsd test.

Where data is only available from the washout period, a linear model was fitted with the measure as the response, and diet group as the predictor variable. The difference between the test diet groups and the control diet were calculated using a Tukey test.

All analyses were performed using R version 3.6.1^(21)^ with the lme4^(22)^ and multcomp^(23)^ libraries. Multiple comparisons correction was applied within each measure to maintain a false discovery rate of 5%.

## Results

### Body weight and intakes

To ensure that the diets had been accepted by the cats for the duration of the study, body weight and intake were investigated. Body weight did not significantly change between diet groups or over the course of the study (Table 2), however intake reduced between baseline and 27 weeks across all groups. Compared to control diet, mean exposure to P and Ca was greater in the test diets (Table 2) reflecting the differences in formulations.

**Table 2.** Body weight, intakes and estimated P and Ca exposure at beginning and end of the study.

|  |  | Control | Moderate | High |
| --- | --- | --- | --- | --- |
|  | baseline (n=75) | 27 weeks (n=25) | 27 weeks (n=25) | 27 weeks (n=25) |
| Mean Body weight | 4.80 | 4.82 | 4.91 | 4.88 |
| (kg: min, max) | (3.01, 7.53) | (2.98, 7.71) | (3.07, 8.10) | (3.59, 6.33) |
| Mean Intake | 62.79 | 52.98 | 57.05 | 59.16 |
| (g/d: min, max) | (35.86, 83.90) | (31.64, 78.68) | (36.28, 75.36) | (42.64, 80.30) |
| Mean P exposure | 0.08 | 0.07 | 0.20 | 0.25 |
| (g/kg MBW/d: min, max) | (0.03, 0.11) | (0.05, 0.10) | (0.15, 0.25) | (0.19, 0.32) |
| Mean Ca exposure | 0.07 | 0.06 | 0.18 | 0.29 |
| (g/kg MBW/d: min, max) | (0.01, 0.10) | (0.02, 0.08) | (0.11, 0.25) | (0.18, 0.39) |

### Mineral apparent digestibility, balance and fractional excretion

The digestibility of P and Ca in the diets and the effect on mineral balance and fractional excretion were evaluated. At baseline, P and Ca excretion and balance were not significantly different between groups (Figure 1). Excretion of P in urine and faeces increased significantly in cats fed both test diets (p<0.001, Figure 1a and c). Urinary excretion of P was higher in cats fed the moderate test diet than the high test diet (p=0.033, Figure 1a), whilst P faecal excretion was higher in cats fed the high test diet than the moderate test diet (p<0.001, Figure 1c). This resulted in a higher positive P balance in cats fed the high test diet than the moderate test diet at the end of the study (p=0.043, Figure 1e). Urinary Ca excretion decreased from baseline in all diet groups (p<0.001, Figure 1b). Faecal excretion of Ca increased after feeding both test diets, although this was significantly greater for the high test diet compared to the moderate test diet (p<0.001, Figure 1d). This resulted in a higher positive Ca balance in cats fed the high test diet than the moderate test diet at the end of the study (p=0.002, Figure 1f).

**Figure 1.**
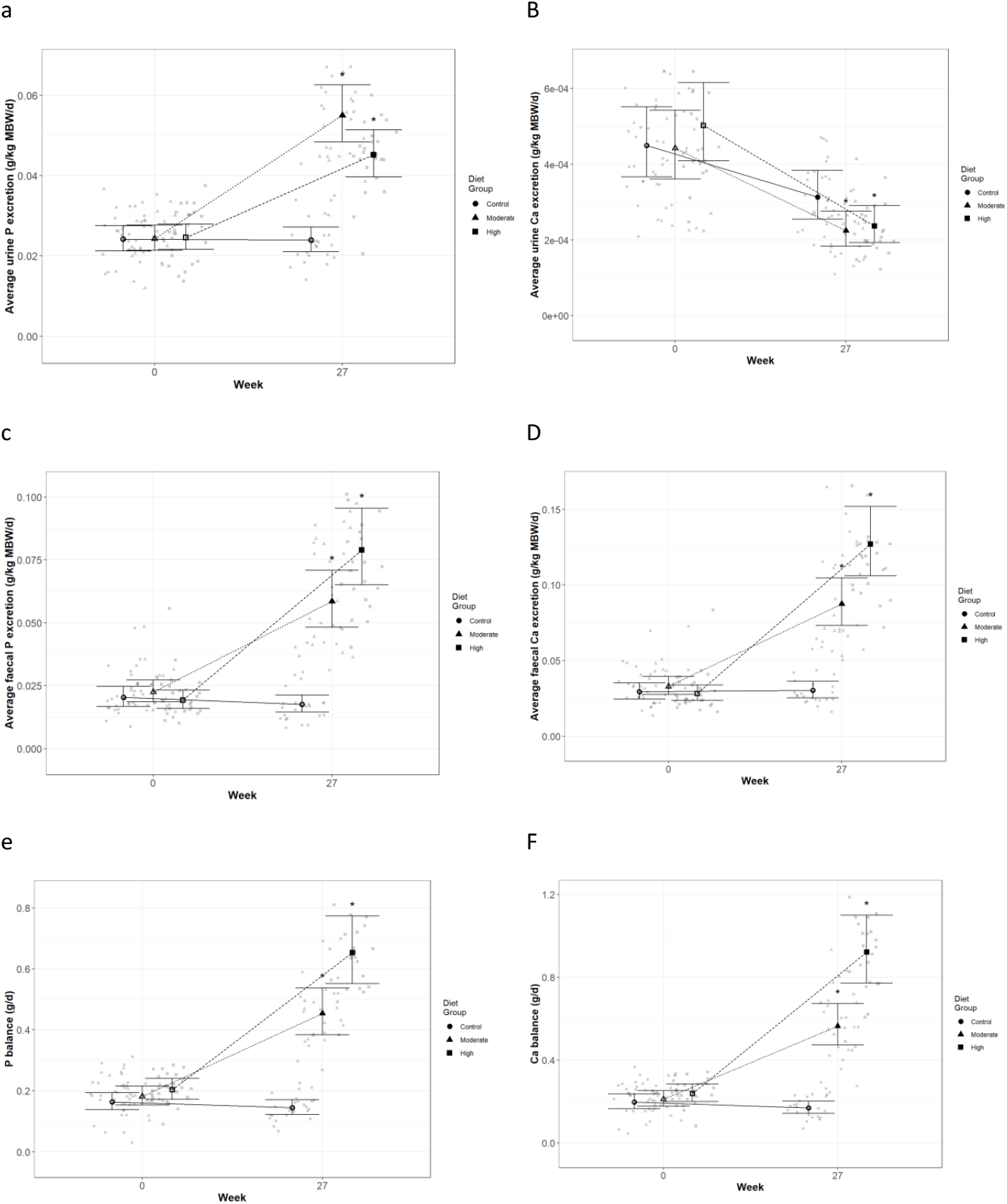
Mean values with 95% upper and lower confidence intervals (n=25 per diet group) for calcium (Ca) and phosphorus (P) excretion and balance at baseline (week 0) and week 27. Urinary excretion (g/kg metabolic body weight, MBW) of a) P and b) Ca; faecal excretion (g/kg MBW) of c) P and d) Ca; and resulting balance (g/d) for e) P and f) Ca in cats fed ● Control (1.4g total P, Ca:P = 0.96), ▲ Moderate (4g total P, 1g P from StPP, Ca:P = 1), and ■ High (5g total P, 1g P from STPP, Ca:P = 1.3) diets. All graphs show baseline and week 27 and are shown by dietary group. * signifies that the change from baseline in that test diet is significantly different from the change from baseline in the control diet, whilst a marker change from an “open” to a “filled” symbol denotes a significant difference from baseline within diet group.

Data indicate fractional excretion of P and Ca did not change for cats fed control diet. However, cats fed test diets increased fractional excretion of P (p<0.001) compared to baseline, whilst fractional Ca excretion reduced (p<0.002, table 3). The fractional excretion of P in the moderate diet group was significantly higher than the high diet group at 27 weeks (p<0.001, table 3).

**Table 3.**
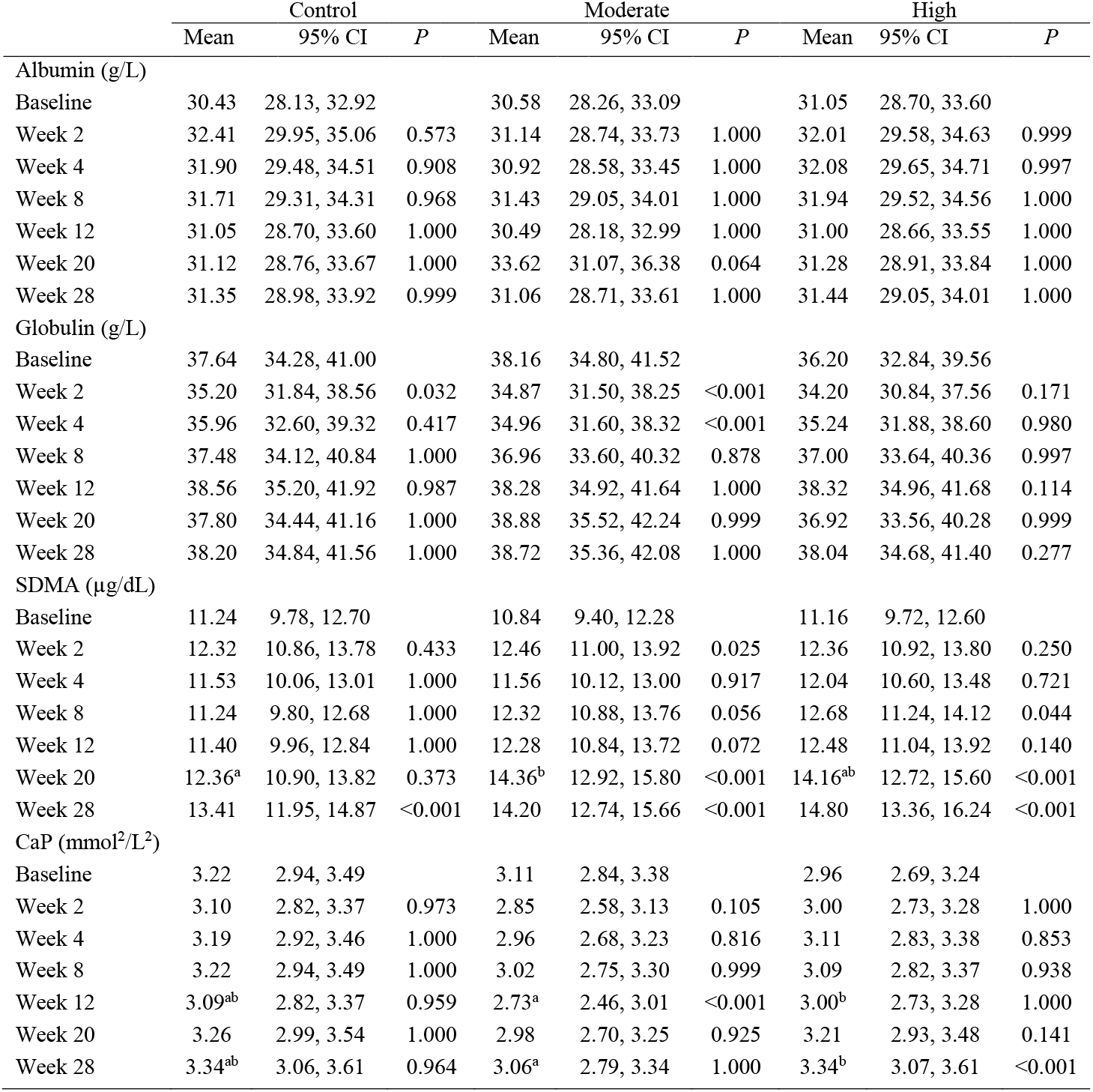
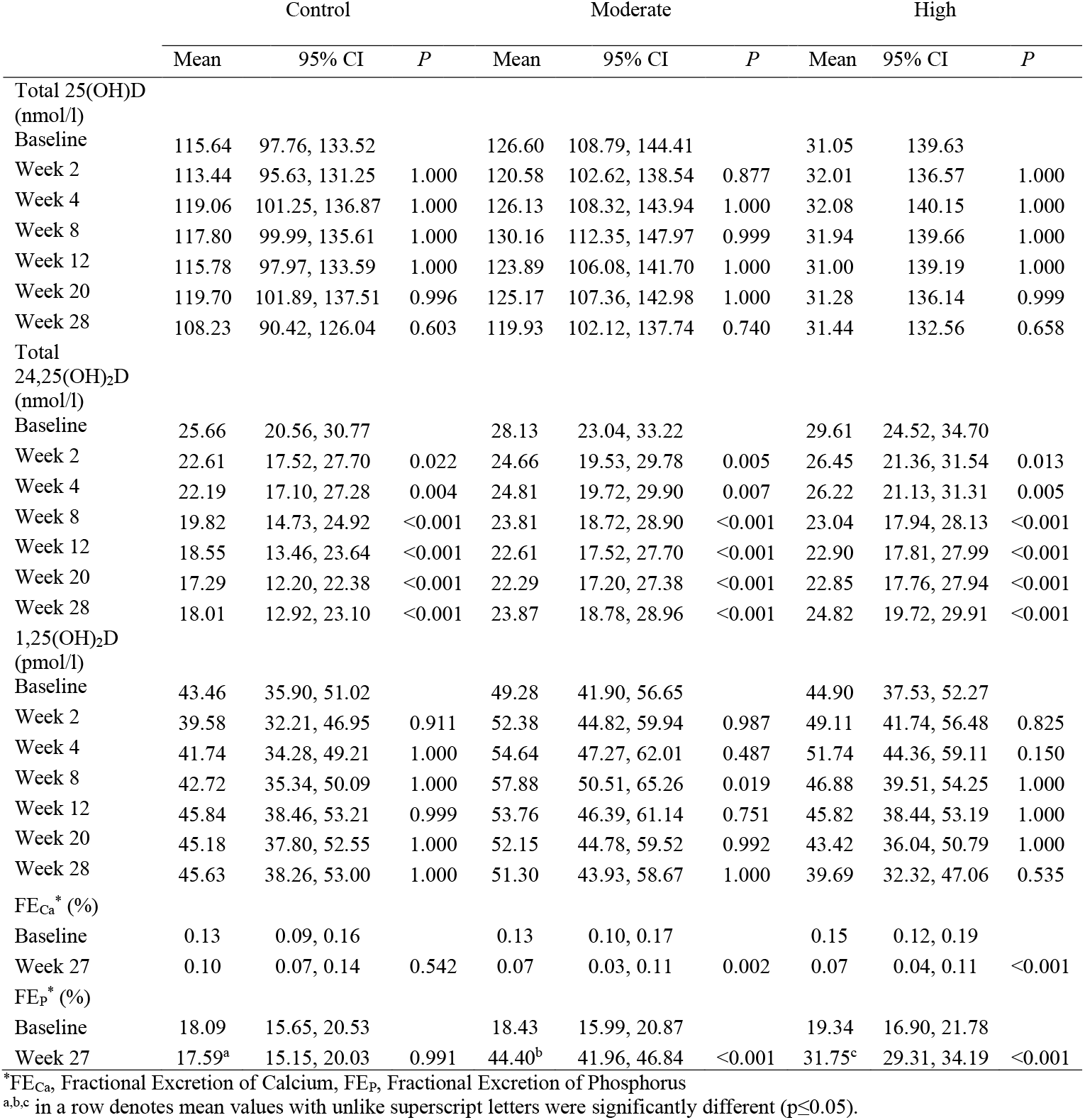
Selected blood biochemistry, vitamin D metabolites and urinary fractional excretion data (mean values and 95% confidence intervals, change from baseline within diet p values reported)

### Relative supersaturation (RSS)

To better understand the risk of renal and urinary stone formation, urine was collected and analysed at the beginning and end (week 0 and 27) of the study for RSS. The RSS for calcium oxalate (CaOX; Figure 2a) was reduced compared to baseline in cats fed both test diets for 27 weeks (p<0.001). At the end of the study, cats fed the high test diet had reduced CaOX RSS compared to those fed the control diet (p=0.005). In contrast, struvite; magnesium ammonium phosphate (MAP), RSS (Figure 2b) was increased cats fed all three experimental diets (p≤0.044) compared to baseline. Cats fed the high test diet had increased MAP compared to cats fed the moderate test diet (p<0.001). Coincidentally, cats fed the high test diet had lower MAP at baseline compared to cats fed the moderate test diet (p=0.005).

**Figure 2.**
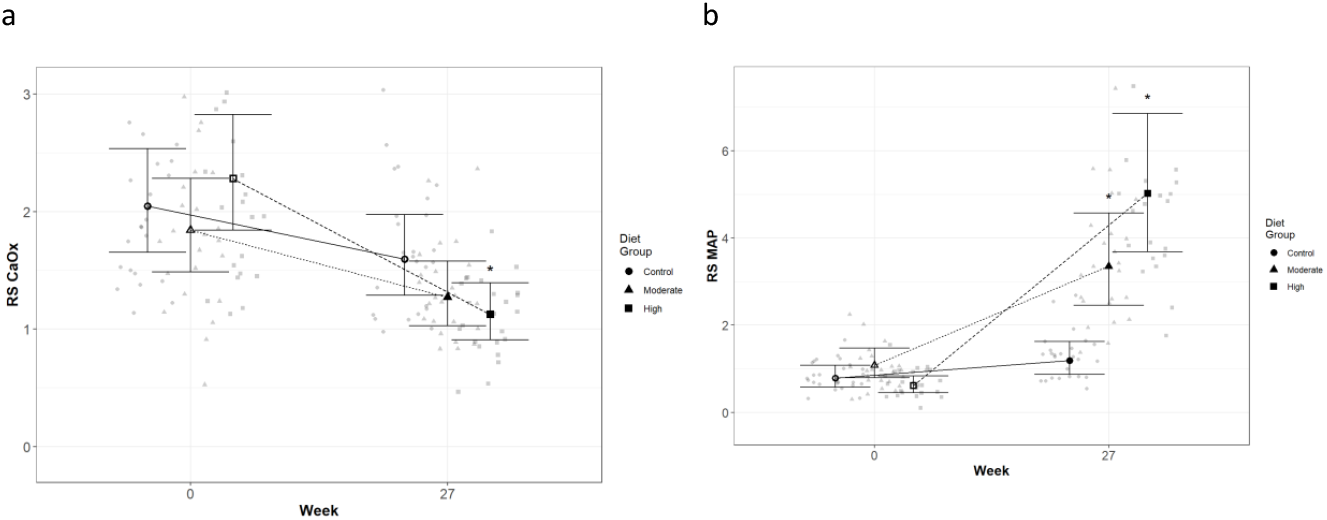
Mean values and 95% upper and lower confidence intervals for urinary relative super saturation of a) calcium oxalate (CaOX) and b) magnesium ammonium phosphate (MAP) at baseline (week 0) and week 27 for cats fed diets ● Control (1.4g total P, Ca:P = 0.96), ▲ Moderate (4g total P, 1g P from STPP, Ca:P = 1), and ■ High (5g total P, 1g P from STPP, Ca:P = 1.3) diets. All graphs show baseline and week 27 and are shown by dietary group. * signifies that the change from baseline in that test diet is significantly different from the change from baseline in the control diet, whilst a marker change from an “open” to a “filled” symbol denotes a significant difference from baseline within diet group.

### Glomerular filtration rate (GFR)

In order to directly assess kidney function, iohexol clearance was assessed at regular intervals to determine GFR. Although no significant differences between diets were detected (Figure 3), a slight decrease from baseline, which did not meet the level of statistical significance for the high diet group, was observed over the course of the study. At week 20, a significant reduction from baseline (p=0.011) was observed for cats fed the control diet, which was not detected at week 28. Cats fed the moderate test diet had significantly lower GFR at week 28 than at baseline (p<0.001), but no difference between diet groups were detected at this time point.

**Figure 3.**
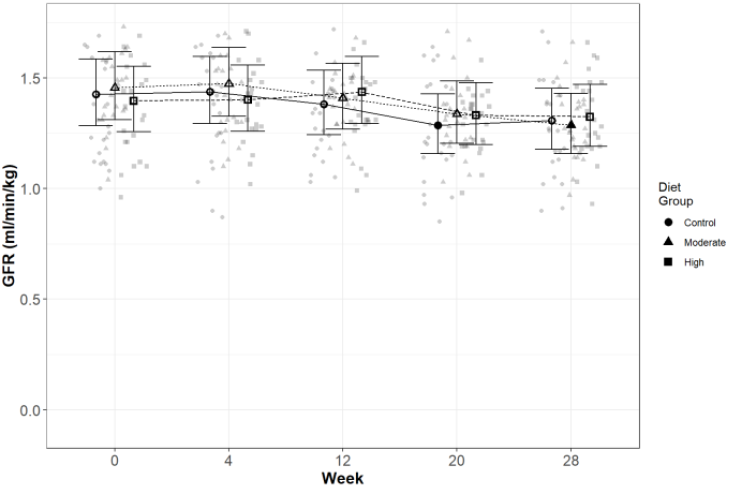
Mean values and 95% upper and lower confidence intervals for glomerular filtration rate (iohexol clearance in ml/min/kg) at baseline (week 0) and over the course of 28 weeks for cats fed diets ● Control (1.4g total P, Ca:P = 0.96), ▲ Moderate (4g total P, 1g P from STPP, Ca:P = 1), and ■ High (5g total P, 1g P from STPP, Ca:P = 1.3) diets. All graphs show baseline and week 27 and are shown by dietary group. * signifies that the change from baseline in that test diet is significantly different from the change from baseline in the control diet, whilst a marker change from an “open” to a “filled” symbol denotes a significant difference from baseline within diet group.

### Blood biochemistry, mineral concentrations, regulatory hormones and bone markers

The blood parameters used as indices of health, kidney and bone function, as well as mineral metabolism and regulation are provided in Figures 4–5 and Tables 3–4.

**Figure 4.**
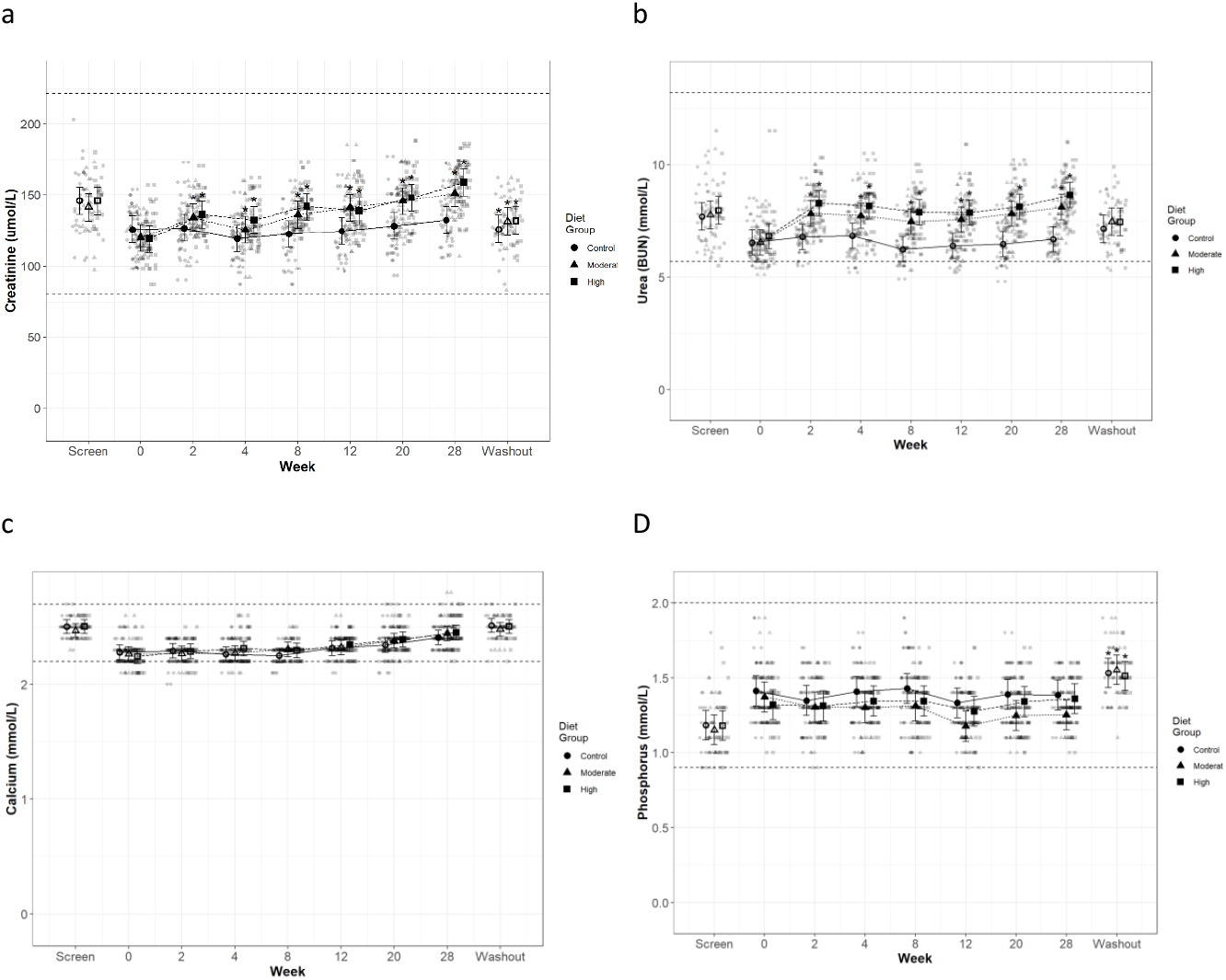
Mean values and 95% upper and lower confidence intervals at baseline (week 0) and during the course of 28 days of feeding for serum biochemical parameters a) creatinine (umol/l), b) urea (mmol/l), c) total calcium (mmol/l) and d) inorganic phosphorus (mmol/l) in cats fed diets ● Control (1.4g total P, Ca:P = 0.96), ▲ Moderate (4g total P, 1g P from STPP, Ca:P = 1), and ■ High (5g total P, 1g P from STPP, Ca:P = 1.3) diets. All graphs show baseline and week 27 and are shown by dietary group. * signifies that the change from baseline in that test diet is significantly different from the change from baseline in the control diet, whilst a marker change from an “open” to a “filled” symbol denotes a significant difference from baseline within diet group. Dotted lines across the graphs indicate the physiological reference ranges from IDEXX.

**Table 4.**
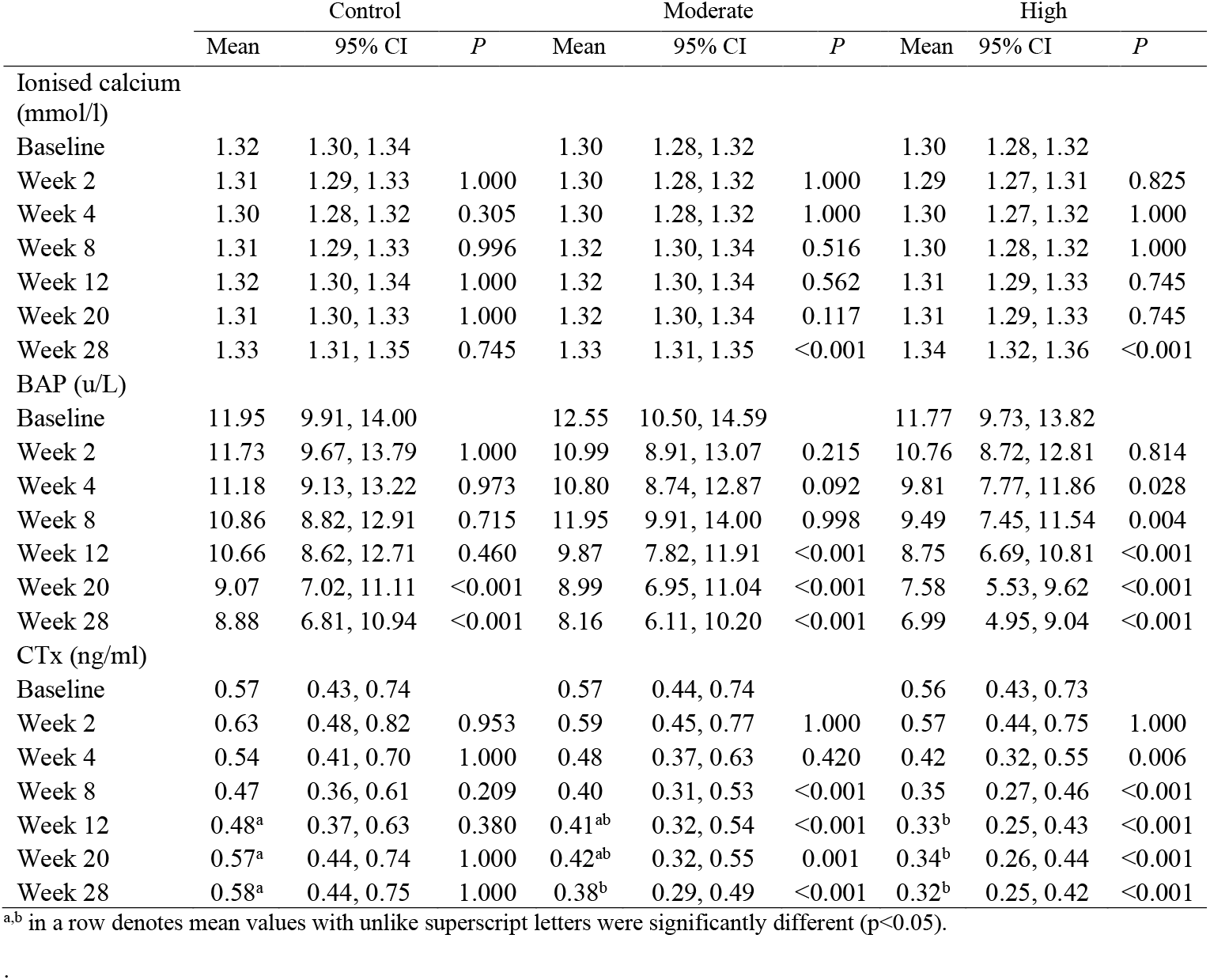
iCa and bone markers (mean values and 95% confidence intervals, change from baseline within diet p values reported)

|  | Control |  |  | Moderate |  |  | High |  |  |
| --- | --- | --- | --- | --- | --- | --- | --- | --- | --- |
|  | Mean | 95% CI | *P* | Mean | 95% CI | *P* | Mean | 95% CI | *P* |
| Ionised calcium (mmol/l) |  |  |  |  |  |  |  |  |  |
| Baseline | 1.32 | 1.30, 1.34 |  | 1.30 | 1.28, 1.32 |  | 1.30 | 1.28, 1.32 |  |
| Week 2 | 1.31 | 1.29, 1.33 | 1.000 | 1.30 | 1.28, 1.32 | 1.000 | 1.29 | 1.27, 1.31 | 0.825 |
| Week 4 | 1.30 | 1.28, 1.32 | 0.305 | 1.30 | 1.28, 1.32 | 1.000 | 1.30 | 1.27, 1.32 | 1.000 |
| Week 8 | 1.31 | 1.29, 1.33 | 0.996 | 1.32 | 1.30, 1.34 | 0.516 | 1.30 | 1.28, 1.32 | 1.000 |
| Week 12 | 1.32 | 1.30, 1.34 | 1.000 | 1.32 | 1.30, 1.34 | 0.562 | 1.31 | 1.29, 1.33 | 0.745 |
| Week 20 | 1.31 | 1.30, 1.33 | 1.000 | 1.32 | 1.30, 1.34 | 0.117 | 1.31 | 1.29, 1.33 | 0.745 |
| Week 28 | 1.33 | 1.31, 1.35 | 0.745 | 1.33 | 1.31, 1.35 | <0.001 | 1.34 | 1.32, 1.36 | <0.001 |
| BAP (u/L) |  |  |  |  |  |  |  |  |  |
| Baseline | 11.95 | 9.91, 14.00 |  | 12.55 | 10.50, 14.59 |  | 11.77 | 9.73, 13.82 |  |
| Week 2 | 11.73 | 9.67, 13.79 | 1.000 | 10.99 | 8.91, 13.07 | 0.215 | 10.76 | 8.72, 12.81 | 0.814 |
| Week 4 | 11.18 | 9.13, 13.22 | 0.973 | 10.80 | 8.74, 12.87 | 0.092 | 9.81 | 7.77, 11.86 | 0.028 |
| Week 8 | 10.86 | 8.82, 12.91 | 0.715 | 11.95 | 9.91, 14.00 | 0.998 | 9.49 | 7.45, 11.54 | 0.004 |
| Week 12 | 10.66 | 8.62, 12.71 | 0.460 | 9.87 | 7.82, 11.91 | <0.001 | 8.75 | 6.69, 10.81 | <0.001 |
| Week 20 | 9.07 | 7.02, 11.11 | <0.001 | 8.99 | 6.95, 11.04 | <0.001 | 7.58 | 5.53, 9.62 | <0.001 |
| Week 28 | 8.88 | 6.81, 10.94 | <0.001 | 8.16 | 6.11, 10.20 | <0.001 | 6.99 | 4.95, 9.04 | <0.001 |
| CTx (ng/ml) |  |  |  |  |  |  |  |  |  |
| Baseline | 0.57 | 0.43, 0.74 |  | 0.57 | 0.44, 0.74 |  | 0.56 | 0.43, 0.73 |  |
| Week 2 | 0.63 | 0.48, 0.82 | 0.953 | 0.59 | 0.45, 0.77 | 1.000 | 0.57 | 0.44, 0.75 | 1.000 |
| Week 4 | 0.54 | 0.41, 0.70 | 1.000 | 0.48 | 0.37, 0.63 | 0.420 | 0.42 | 0.32, 0.55 | 0.006 |
| Week 8 | 0.47 | 0.36, 0.61 | 0.209 | 0.40 | 0.31, 0.53 | <0.001 | 0.35 | 0.27, 0.46 | <0.001 |
| Week 12 | 0.48<sup>a</sup> | 0.37, 0.63 | 0.380 | 0.41<sup>ab</sup> | 0.32, 0.54 | <0.001 | 0.33<sup>b</sup> | 0.25, 0.43 | <0.001 |
| Week 20 | 0.57<sup>a</sup> | 0.44, 0.74 | 1.000 | 0.42<sup>ab</sup> | 0.32, 0.55 | 0.001 | 0.34<sup>b</sup> | 0.26, 0.44 | <0.001 |
| Week 28 | 0.58<sup>a</sup> | 0.44, 0.75 | 1.000 | 0.38<sup>b</sup> | 0.29, 0.49 | <0.001 | 0.32<sup>b</sup> | 0.25, 0.42 | <0.001 |

<sup>a,b</sup> in a row denotes mean values with unlike superscript letters were significantly different (p<0.05).

Secondary markers of kidney health, creatinine (Figure 4a) and blood urea nitrogen (BUN; Figure 4b), were increased in cats fed both test diets when compared to baseline values and to those fed the control diet (p≤0.041 and p<0.001, respectively). Serum SDMA (Table 3) increased in all groups over the course of the study (p<0.001) and change from baseline for cats fed the moderate test diet was significantly greater than that for cats fed the control diet at week 20 (p=0.015). In contrast, no difference between the control and the high test diet group were detected. Serum albumin and globulin concentrations (Table 3) remained unchanged throughout the study with no effect of diet observed, although a reduction between screening and baseline was detected for cats fed the control diet and the high test diet (serum albumin p<0.001) and cats fed the high test diet only (serum globulin p=0.004). Serum albumin concentration was found to significantly decrease between screening and washout period for cats fed the high test diet (p=0.03, Table 3).

Total serum Ca (Figure 4c) increased in all diet groups by the end of the study, from week 12 for the high test diet, week 20 for the moderate test diet and week 28 for the control diet(p<0.001). Serum inorganic P (Figure 4d) remained stable throughout the study, except in cats fed the moderate test diet where P was decreased at weeks 12 to 28 when compared to baseline values (p≤0.005). There were, however, no significant between diet differences in serum P concentration during the study. From baseline to week 28, cats fed the high test diet significantly increased CaP product concentration (Table 3) (p<0.001), but there was no significant difference compared to the control group at any time. However, at weeks 12 and 28 the change from baseline in the moderate test diet group was significantly less than that in the high test diet group (p=0.03 and 0.02 respectively). There was a significant increase in CaP product observed for all diets when screening and washout periods were compared (p<0.001, Table 3).

The plasma concentration of regulatory hormone PTH (Figure 5a) did not show a significant effect of diet, but did decline over time for all diet groups (p≤0.01). However, FGF-23 plasma concentrations (Figure 5b) were significantly increased in cats fed the moderate test diet (p≤0.028). At week 12 cats fed the high test diet had increased FGF-23 concentrations (Figure 5b, p<0.001), this was returned to baseline at week 20, and cats fed the control diet showed a significant reduction in FGF-23 at week 28 (Figure 5b, p=0.012).

**Figure 5.**
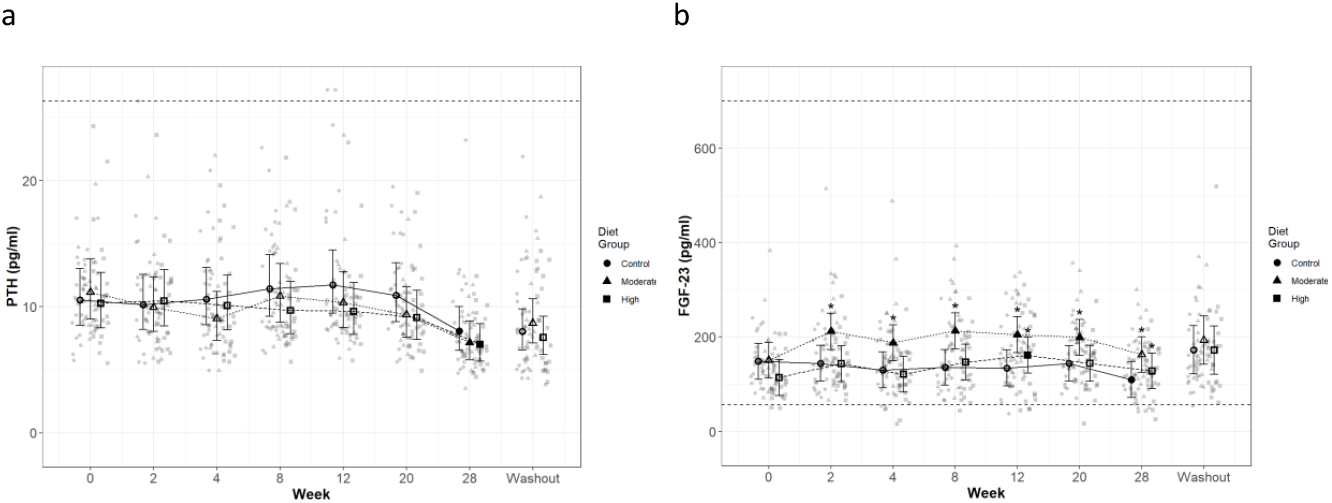
Mean values and 95% upper and lower confidence intervals at baseline (week 0) and during the course of 28 days of feeding for plasma concentrations of a) parathyroid hormone (PTH, pg/ml) and b) fibroblast growth factor – 23 (FGF-23, pg/ml) in cats fed ● Control (1.4g total P, Ca:P = 0.96), ▲ Moderate (4g total P, 1g P from STPP, Ca:P = 1), and ■ High (5g total P, 1g P from STPP, Ca:P = 1.3) diets. All graphs show baseline and week 27 and are shown by dietary group. * signifies that the change from baseline in that test diet is significantly different from the change from baseline in the control diet, whilst a marker change from an “open” to a “filled” symbol denotes a significant difference from baseline within diet group. Dotted lines across the graphs indicate the physiological reference ranges.

Vitamin D metabolites (Table 3) total 25(OH)D, total 24,25(OH)_2_D and 1,25(OH)_2_D showed no significant differences in serum concentration between the diet groups for the duration of the study (p>0.05). Total 24,25(OH)_2_D concentrations decreased significantly from baseline in all diet groups from week 2 (control p=0.022, moderate test diet p=0.005 and high test diet p=0.013). Cats fed the high test diet showed a significant increase from baseline in 1,25(OH)_2_D concentration at week 8 (p=0.019), then returned to baseline levels for the rest of the study.

Among the markers of bone turnover measured, blood ionised calcium (iCa) concentrations in cats fed the test diets were significantly increased from baseline levels at week 28 (p<0.001, Table 4). Bone alkaline phosphatase (BAP) concentrations were not significantly different between diet groups, but decreased from baseline over time for all diet groups (Table 4). This decrease was, however, observed earlier in cats fed the high test diet, which was significantly reduced from baseline from week 4 onward (p=0.028). At week 12, BAP concentrations for cats fed the moderate test diet were also significantly reduced compared to baseline (p<0.001), while for the control diet group a significant reduction was detected from week 20 (p<0.001). For serum cross-laps (CTx), no change from baseline was observed over the course of the study for the control diet group (Table 4.0). Cats fed test diets, however, showed a decrease in CTx concentrations over time (p≤0.006, Table 4). This decrease from baseline was significantly greater than the control group from week 12 for cats fed the high test diet and week 28 for the moderate test diet (p=0.03 and p<0.001, respectively). After the washout period cats fed control diet had significantly lower CTx than at week 28 (p<0.001).

### Haematology

Haematology analysis was carried out as a measure of health, those parameters affected by diet are reported in Table 5. Mean corpuscular haemoglobin (MCH), mean corpuscular volume (MCV) and reticulocyte haemoglobin reduced significantly over the course of the study for all diet groups, whilst staying within IDEXX laboratory reference ranges (p<0.001). At weeks 12 and 28, cats fed the moderate and high test diets were observed to have significantly decreased MCH, compared to the control diet group, (p=0.04 and p=0.01 respectively). MCV for cats fed test diets was also observed to have reduced significantly more than the control diet group at week 12 (moderate test diet p=0.002, high test diet p=0.024) and this remained significantly lower than the control group at week 20 in cats fed the high test diet (p=0.04). MCH and MCV were also found to significantly decrease between screening and washout periods for all diet groups (p<0.001). Compared to cats fed control diet, the MCH of cats fed the moderate test diet was significantly decreased (p=0.023) and the MCV of cats fed the high test diet was significantly decreased (p=0.029) following the washout period. The concentration of reticulocyte haemoglobin in cats fed the moderate test diet was significantly reduced compared to the control diet group from week 12 onwards (p≤0.042).

**Table 5.** Selected haematology (mean values and 95% confidence intervals change from baseline within diet p values reported)

|  | Control |  |  | Moderate |  |  | High |  |  |
| --- | --- | --- | --- | --- | --- | --- | --- | --- | --- |
|  | Mean | 95% CI | *P* | Mean | 95% CI | *P* | Mean | 95% CI | *P* |
| Mean Corpuscular Haemoglobin (pg) |  |  |  |  |  |  |  |  |  |
| Baseline | 14.96 | 14.44, 15.48 |  | 15.01 | 14.49, 15.52 |  | 15.01 | 14.50, 15.53 |  |
| Week 2 | 14.83 | 14.31, 15.34 | 0.999 | 14.95 | 14.43, 15.47 | 1.000 | 14.93 | 14.41, 15.45 | 1.000 |
| Week 4 | 14.72 | 14.20, 15.24 | 0.748 | 14.61 | 14.09, 15.12 | 0.074 | 14.55 | 14.03, 15.06 | 0.016 |
| Week 8 | 14.36 | 13.85, 14.88 | <0.001 | 13.97 | 13.46, 14.49 | <0.001 | 14.00 | 13.48, 14.52 | <0.001 |
| Week 12 | 14.00<sup>a</sup> | 13.49, 14.52 | <0.001 | 13.52<sup>ab</sup> | 13.00, 14.03 | <0.001 | 13.45<sup>b</sup> | 12.93, 13.96 | <0.001 |
| Week 20 | 14.01 | 13.49, 14.52 | <0.001 | 13.48 | 12.96, 14.00 | <0.001 | 13.50 | 12.98, 14.02 | <0.001 |
| Week 28 | 13.72<sup>a</sup> | 13.21, 14.24 | <0.001 | 13.11<sup>b</sup> | 12.60, 13.63 | <0.001 | 13.23<sup>ab</sup> | 12.71, 13.74 | <0.001 |
| Mean Corpuscular Volume (fl) |  |  |  |  |  |  |  |  |  |
| Baseline | 44.79 | 42.72, 46.86 |  | 44.88 | 42.81, 46.95 |  | 44.40 | 42.33, 46.48 |  |
| Week 2 | 45.04 | 42.96, 47.11 | 1.000 | 45.24 | 43.16, 47.32 | 1.000 | 44.67 | 42.59, 46.75 | 1.000 |
| Week 4 | 44.80 | 42.72, 46.88 | 1.000 | 44.21 | 42.14, 46.28 | 0.903 | 43.96 | 41.89, 46.04 | 0.998 |
| Week 8 | 43.96 | 41.89, 46.03 | 0.659 | 42.49 | 40.42, 44.56 | <0.001 | 41.94 | 39.86, 44.01 | <0.001 |
| Week 12 | 43.24<sup>a</sup> | 41.17, 45.31 | 0.010 | 40.86<sup>b</sup> | 38.79, 42.93 | <0.001 | 40.81<sup>b</sup> | 38.74, 42.88 | <0.001 |
| Week 20 | 42.14<sup>a</sup> | 40.06, 44.21 | <0.001 | 40.37 | 38.30, 42.44 | <0.001 | 39.81<sup>b</sup> | 37.74, 41.88 | <0.001 |
| Week 28 | 41.22 | 39.15, 43.29 | <0.001 | 39.60 | 37.52, 41.67 | <0.001 | 39.80 | 37.72, 41.87 | <0.001 |
| Reticulocyte Haemoglobin (pg) |  |  |  |  |  |  |  |  |  |
| Baseline | 16.21 | 15.51, 16.90 |  | 16.83 | 16.13, 17.52 |  | 16.44 | 15.74, 17.14 |  |
| Week 2 | 16.27 | 15.57, 16.96 | 1.000 | 16.33 | 15.63, 17.04 | 0.525 | 16.04 | 15.34, 16.75 | 0.829 |
| Week 4 | 15.93 | 15.22, 16.63 | 0.990 | 15.82 | 15.13, 16.52 | <0.001 | 15.91 | 15.21, 16.60 | 0.387 |
| Week 8 | 15.66 | 14.96, 16.35 | 0.330 | 15.62 | 14.92, 16.31 | <0.001 | 15.16 | 14.46, 15.86 | <0.001 |
| Week 12 | 15.52<sup>a</sup> | 14.83, 16.22 | 0.088 | 15.00<sup>b</sup> | 14.30, 15.70 | <0.001 | 15.06<sup>ab</sup> | 14.36, 15.76 | <0.001 |
| Week 20 | 15.34<sup>a</sup> | 14.64, 16.04 | 0.007 | 14.91<sup>b</sup> | 14.21, 15.60 | <0.001 | 14.75<sup>ab</sup> | 14.05, 15.44 | <0.001 |
| Week 28 | 15.09<sup>a</sup> | 14.40, 15.79 | <0.001 | 14.63<sup>b</sup> | 13.93, 15.32 | <0.001 | 14.93<sup>ab</sup> | 14.24, 15.63 | <0.001 |

<sup>a,b</sup> in a row denotes mean values with unlike superscript letters were significantly different (p<0.05).

### Urinalysis

To assess the general health of the kidneys and urinary tract, urinalysis was performed throughout the study. Urine pH increased in cats fed the moderate (p≤0.02) and high test diets (p<0.001), compared to baseline, from weeks 8 and 2 respectively (Figure 6a). This measure was also significantly increased for cats fed the high test diet, compared to cats fed control diet, at week 2 (p=0.017) and from week 12 onwards (p≤0.012). Urine pH in cats fed the moderate test diet was significantly higher than in the control group only at week 12 (p=0.045). Urine specific gravity was observed to significantly increase in cats fed the moderate test diet from weeks 4 to 20 (p≤0.07). In cats fed the high test diet, urine specific gravity was observed to be increased between weeks 4 and 12 (p≤0.016) before returning to baseline levels at week 20 (Figure 6b). Despite oscillations over the study, urine protein:creatinine ratio (UPCR) was reduced compared to baseline in all cats regardless of diet group (control diet, week 27 p=0.002; moderate test diet, week 12 p=0.015; and high test diet, week 20 p=0.014; Figure 6c). No significant effect of time or diet on the urine albumin:creatinine ratio (uACR) was observed (Figure 6d).

**Figure 6.**
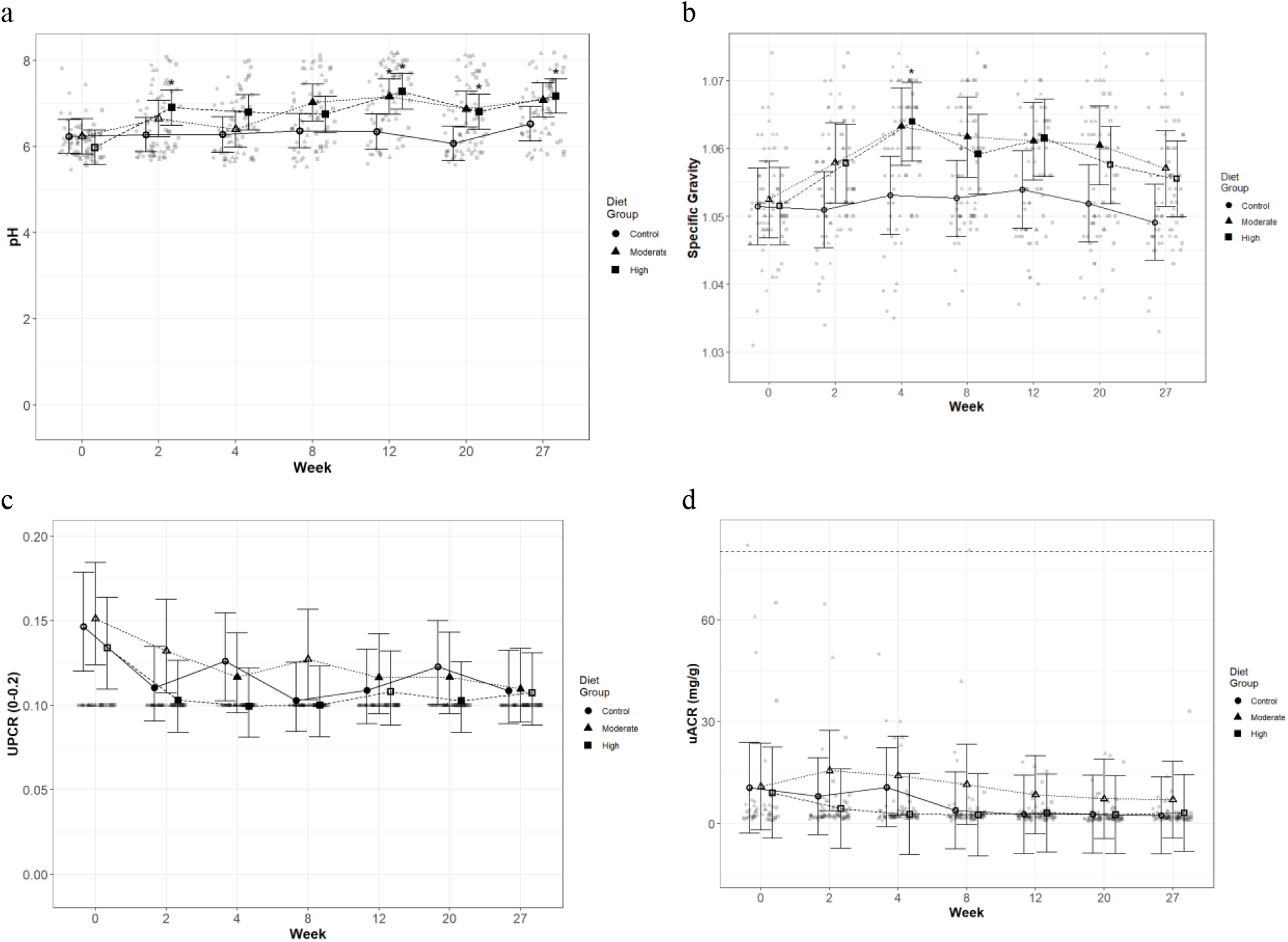
Mean values and 95% upper and lower confidence intervals for urine parameters at baseline (week 0) and during the course of 27 weeks for a) pH, b) urine specific gravity, c) protein:creatinine ratio (UPCR), and d) microalbumin:creatinine ratio (uACR) for cats fed diets ● Control (1.4g total P, Ca:P = 0.96), ▲ Moderate (4g total P, 1g P from STPP, Ca:P = 1), and ■ High (5g total P, 1g P from STPP, Ca:P = 1.3) diets. All graphs show baseline and week 27 and are shown by dietary group. * signifies that the change from baseline in that test diet is significantly different from the change from baseline in the control diet, whilst a marker change from an “open” to a “filled” symbol denotes a significant difference from baseline within diet group. Dotted lines across UACR graph indicate the upper physiological reference range

### Abdominal ultrasound and DXA

In order to assess soft tissue and skeletal mineralisation, imaging via DXA and ultrasound was performed. No changes in mineralisation were observed during DXA scans at baseline or week 28 and no difference in bone mineral density (BMD) were detected following 28 weeks of feeding the experimental diets (Figure 7).

**Figure 7.**
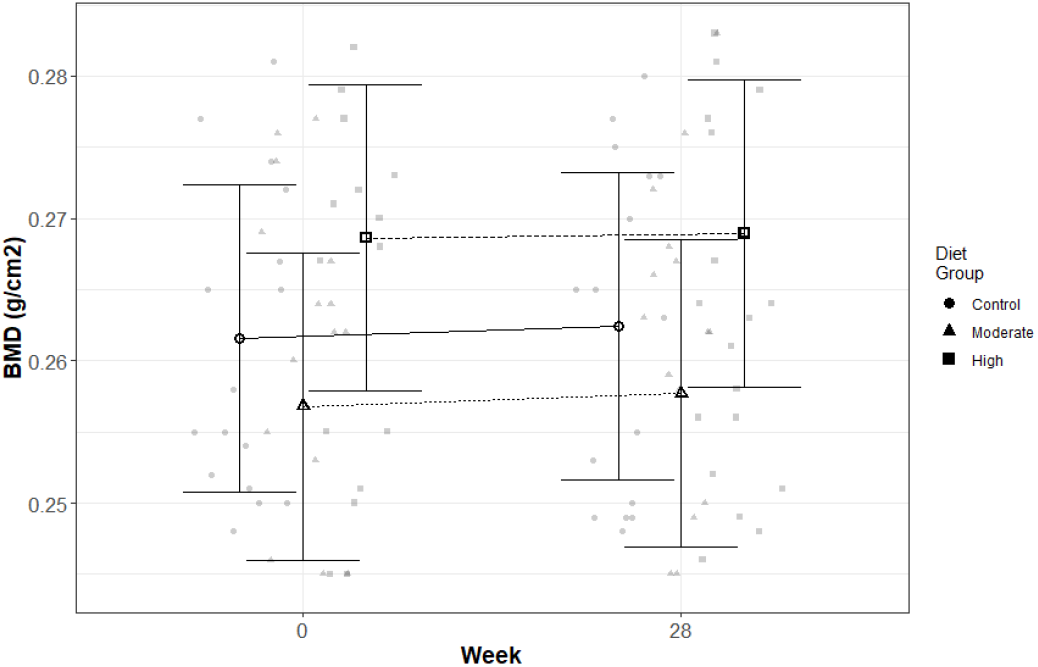
Mean values and 95% upper and lower confidence intervals for Bone Mineral Density (BMD; in g/cm^2^) as assessed by DXA at baseline (week 0) and week 28 for cats fed diets ● Control (1.4g total P, Ca:P = 0.96), ▲ Moderate (4g total P, 1g P from STPP, Ca:P = 1), and ■ High (5g total P, 1g P from STPP, Ca:P = 1.3) diets. All graphs show baseline and week 27 and are shown by dietary group. * signifies that the change from baseline in that test diet is significantly different from the change from baseline in the control diet, whilst a marker change from an “open” to a “filled” symbol denotes a significant difference from baseline within diet group.

No changes in kidney size, shape or soft tissue structure indicative of pathology were observed by renal sonography. At the end of the 30 week study, one cat from the moderate test diet group was reported by the primary imager to have partial (or non-continuous) rim signs that were not present at baseline, indicating presence of some echogenicity of the thin medullary band parallel to the corticomedullary junction where the proximal renal tubules reside. Several additional findings were noted in both the first and second opinions, and therefore a frequency analysis was performed (Table 6). This showed that the incidence of rim signs and bright (hyperechoic) renal cortices did not differ between diets (p>0.88). However, it was noted that these findings increased as time progressed in all diet groups. There was also one instance of “speckling” and one cat developed renal “cysts”, but as these were isolated observations, no statistical analyses could be performed.

**Table 6.** Number of cats showing findings in their ultrasound data by diet group and time point. There were 25 cats in each diet group.

|  | Control |  |  | Moderate |  |  | High |  |  | *P* |
| --- | --- | --- | --- | --- | --- | --- | --- | --- | --- | --- |
|  | Base-line | Mid | Final | Base-line | Mid | Final | Base-line | Mid | Final |  |
| rim signs | 0 | 0 | 2 | 2 | 4 | 7 | 1 | 2 | 3 | 0.89 |
| Bright cortices | 13 | 11 | 14 | 8 | 7 | 10 | 7 | 11 | 12 | 0.88 |
| Speckling | 0 | 0 | 0 | 0 | 0 | 1 | 0 | 0 | 0 | N/A |
| Cysts | 0 | 0 | 1 | 0 | 0 | 0 | 0 | 0 | 0 | N/A |

## Discussion

This 30-week feeding study in healthy adult cats was initiated to investigate the safety of phosphorus-rich, dry extruded diets containing 1.0g/1000kcal of a soluble P salt. Results indicate that the diets with total P levels of 4.0 and 5.0g/1000kcal, of which STPP provided 1.0g/1000kcal, and Ca:P ratios of 1.0 and 1.3, respectively, did not cause adverse health effects. Ultrasound and DXA scans did not reveal adverse structural changes in renal or skeletal tissue. Blood and urinary markers of general health, renal function and bone metabolism remained within physiological reference ranges. Findings from the 4-week washout period indicate that the increased serum creatinine and urea values in cats fed the test diets were adaptations to dietary factors, including but not limited to P levels, rather than indications of kidney damage. Differences between diet groups in mineral status and regulators of Ca and P metabolism were detected. However, the absence of adverse health effects suggest that these regulatory mechanisms were operating within physiological boundaries, and adaptation to the added Ca and P load was apparently successful following 30 weeks of exposure. Therefore, this is the first chronic feeding study to demonstrate a no observed adverse effect level (NOAEL) for a soluble P salt in complete and balanced, dry feline diets; specifically STPP providing 1.0g/1000kcal P in diets containing up to 5.0g/1000kcal total P (with Ca:P 1.0 or 1.3) for a 30week feeding period in healthy, adult cats.

### Measures of kidney and bone function and health

No significant structural changes or renoliths were observed in the kidneys on ultrasound imaging after 30 weeks of feeding STPP at this dietary concentration. Although medullary rim signs were observed in some cats over the length of the study, frequency analysis suggested that these were not diet-related. The observed increase in albumin in the urine in association with renal structural changes reported in previous studies^(2)^ were absent here. Nevertheless, feeding the moderate test diet was associated with increased plasma FGF-23 and reduced fasting plasma phosphate concentrations.

While remaining well within physiological reference ranges, creatinine and urea values increased in the cats fed the P-rich test diets. Although increasing levels of these parameters are considered risk factors for developing CKD, other parameters did not indicate cause for concern: GFR and SDMA (an indirect marker of GFR), did not differ between diet groups. The cats fed the P rich diets continued to concentrate their urine well, as indicated by the higher USG values compared to control, and UPCR and UACR were reduced or unchanged indicative of an absence of glomerular disease. Furthermore, comparison of blood and urine values between the screening, test and washout periods indicate that some of the differences detected between experimental groups during the test period were likely caused by diet formulation differences rather than effects of dietary P levels on kidney function or health. For example, the lower protein level (above AAFCO minimum of 26g/100g dry matter) in the control diet compared to the (wet format) screening and test diets (Table 1) may explain the reduced serum creatinine and urea levels at baseline and the subsequent increases, particularly in urea, in cats transitioned onto the test diets with higher protein^(24)^. Likewise, the higher serum creatinine and urea concentrations observed in test diet fed cats decreased following the 4 week washout period to levels lower or similar to those observed during screening. This may indicate that changes induced by the P-rich test diets were transient and not indicative of impaired kidney function or health.

Data reported in older cats (mean age 10.1 years) in a 2-year study^(25)^, focussed on the effect of sodium content, may provide support for the longer term safety of the P-rich test diets in the current study beyond 7 months. The diets fed by Reynolds *et. al*. in this study contained lower total P and Ca:P, with a mean of 2.3g/1000kcal and Ca:P of 0.8, but were similar to the moderate test diet in inorganic soluble P salt contribution (1g/1000kcal; Vincent Biourge, personal communication). During this 2 year study, no detrimental effects on GFR or structural changes in the kidneys as assessed by ultrasound, or albuminuria were observed. However, Reynolds et al^(25)^ did not report a decrease in serum P concentrations, possibly due to the lower total total P content of the diets and due to the lack of FGF-23 data it is not possible to fully compare the studies to predict safety beyond 30 weeks.

In the current study, DXA scans did not reveal skeletal changes at baseline or after 30 weeks of feeding the test diets, nor did BMD and BAP values differ between diet groups. The serum CTx concentration suggested that bone mineral resorption was reduced in cats fed the Ca and P-rich test diets. Despite no changes detected with DXA scans in control group cats, CTx values indicated that mineral resorption may be occurring, but was reduced following the washout period. The control diet was lower in Ca, P, Ca:P as well as protein than the other two test diets (Table 1), which may have induced an increase in bone turnover although no changes in BMD were detected. Yamka *et. al*.^(26)^ showed that increasing dietary crude protein, calcium and Ca:P content (along with increases in eicosapentaenoic, docosahexaenoic and n-3 fatty acids) reduced CTx in healthy geriatric cats, corroborating findings from the high test diet group, containing the highest protein, Ca and Ca:P ratios of the three experimental diets, had the lowest CTx levels after week 12 of the study.

### Mineral metabolism and regulatory mechanisms

In response to increased total and highly soluble P salt inclusion in the test diets, cats increased P excretion in the urine, as well as the faeces, the latter leading to the reduced apparent P digestibility. However, as the balance data suggest, the digestibility was not low enough to fully compensate for the increased P intake – between 2 and 3-fold higher, respectively, in test compared to control group cats. Total P levels in the test diets dose-dependently increased the positive Ca and P balance in the cats, which differed from findings in the previous studies conducted by our group^(2)^. A test diet containing 3.6g/1000kcal from SDHP (4.8g/1000kcal total P, Ca:P 0.59) resulted in no change in P but a negative Ca balance following 4 weeks of feeding. Decreasing SDHP to 1.5g/1000kcal (3.6g/1000kcal total P, Ca:P 0.93) and feeding for 28 weeks caused a more marked increase in P balance compared that reported here, while no change in Ca balance data was observed^(2)^. These differences between studies appear to be largely driven by the reduction of Ca excreted via the urine in the current study, which was not observed previously. Interestingly, baseline balance values in all three of these studies were positive, supporting the hypothesis that cats tend to have a positive Ca and P balance, even when Ca and P levels in the diet are relatively low, as reported previously^(2; 5; 27)^. Mineral balance is difficult to perform and methodologies used can affect the variation in the results, here overnight losses in urinary volume as noted previously by Hendriks *et al*^(28)^ cannot be accounted for as we did not employ radiolabelling methods. There is also analytical variability in measurement of minerals in the laboratory which may hinder the ability to measure balance with 100% certainty. However, focussing on the positive balances observed for the cats fed the tests diets, detailed insights into regulatory mechanisms and impact on long term health following continuous positive Ca and P burden in cats fed mineral-rich test diets is not fully understood.

To understand the risk of developing soft tissue mineralisation, renoliths or uroliths the CaP product was calculated. In all cats’ CaP product increased during the current study with no adverse effects detected on kidney or bone scans. However, it is important to note that all CaP product values in all diet groups (Table 3) were below the threshold level of 5.6mmol^2^/l^2^ (70mg^2^/dl^2^) at which soft tissue mineralisation has been reported to occur^(29; 30)^, helping to explain the absence of any findings. RSS data suggest that the test group cats may have had a higher risk of struvite (MAP) urolith formation,^(31; 32; 33)^. However, there was no evidence of stone formation in any cats fed the test diets. This is in contrast to data from Alexander *et al*.^(2)^, where no sustained increase in struvite RSS was determined but urolith and renoliths were observed.

The increased phosphorus excretion in faeces and urine in cats fed the test diets indicated homeostatic mechanisms were functioning to enable serum P to be kept relatively stable. While both test diets resulted in an absence of adverse structural or functional effects on the kidney following 30 weeks of feeding, intake of the moderate test diet with the lower Ca:P ratio of 1 resulted in reduced fasting plasma phosphate and elevated FGF-23. At week 8 of the study elevation in calcitriol (1,25(OH)_2_D) levels in cats fed the moderate test diet was also observed, possibly as a result of the increased FGF-23^(34)^, however vitamin D metabolite levels did not otherwise differ between diet groups. These observations may suggest that the moderate P diet activated regulatory mechanisms not required by cats exposed to the high test diet due to its higher Ca:P of 1.3. The small, but significant increases in fasted FGF-23 observed here, cannot be excluded as possible risk factors for the longer term development of renoliths and structural changes, as observed previously^(2)^. However, Alexander *et al*. observed a concurrent increase in PTH, which did not occur in the current study. Both increased PTH and FGF-23 were suggested to contribute to reduced reabsorption of phosphate in the kidney’s proximal tubules, which would explain the increased urinary and fractional P excretion and lower serum P levels, all observed in the current study. This would largely corroborate the current understanding of regulatory mechanisms involved in P homeostasis^(35)^. PTH responses in cats do appear to be more rapid than FGF-23, as indicated by post-prandial profiles observed following acute feeding studies^(7)^, and the PTH measured in fasted blood samples may not indicate those present during the first hours after each meal. Although not measured at screening, interestingly, plasma PTH increased while FGF-23 values did not change between week 28 and the washout period. However other regulatory proteins not measured during this study, such as klotho could influence homeostasis and, as not well understood in cats, merit further investigation.

Calcium and iCa concentrations in blood increased over the course of the study, but did not differ significantly between control or test diets. Feeding the control diet prior to baseline sampling decreased total serum Ca significantly compared to screening values (by ~ 0.2mmol.L), it is not known whether there was a concomitant decrease in iCa as it was not measured at screening. However, all diet groups subsequently increased in iCa concentration during the test period before returning to screening levels. Time-dependent changes in diets (such as water loss etc) or seasonal variability cannot be excluded as factors influencing trends observed across all diet groups.

## 1. Conclusion

The data provide evidence that dry format diets, tested in healthy adult cats, containing 1.0g/1000kcal (4184kJ) P in the form of the soluble salt STPP caused no observable adverse effects when fed for 30 weeks. This was true for both test diets formulated for this study, total P of 4.0 and 5.0g/1000kcal (4184kJ) and Ca:P of 1.0 and 1.3 respectively, additionally providing evidence for NOAEL of a higher inclusion of P from organic sources with an increased Ca:P. Ultrasound and DXA scans did not reveal signs of adverse structural changes in renal or skeletal tissue. Markers of renal function, general and skeletal health, remained within physiological reference ranges. Any differences observed between diet groups were considered regulatory responses to maintain mineral homeostasis or adaptations to dietary factors other than P levels. These findings will assist the pet food industry, regulators and professional bodies in developing guidance on safe maximum dietary levels of P for healthy adult cats.

---

**Supplementary Table –.**
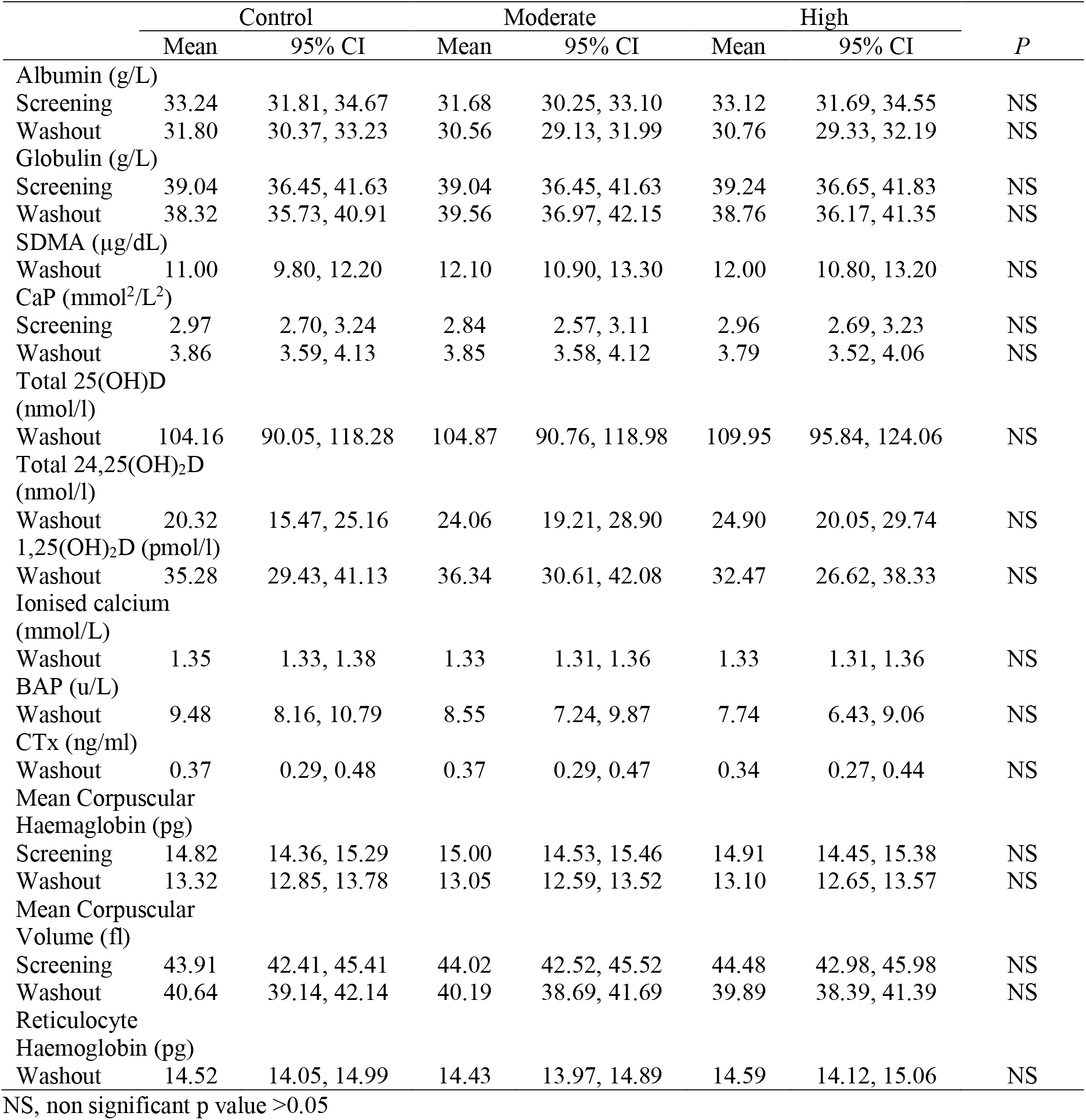
Washout and/or Screening data for tabulated measures found within the main body of the manuscript (mean values and 95% confidence intervals, between diet p values reported)

## References

1. Brown CA, Elliott J, Schmiedt CW et al. (2016) Chronic Kidney Disease in Aged Cats: Clinical Features, Morphology, and Proposed Pathogeneses. Veterinary pathology 53, 309–326.

2. Alexander J, Stockman J, Atwal J et al. (2019) Effects of the long-term feeding of diets enriched with inorganic phosphorus on the adult feline kidney and phosphorus metabolism. British Journal of Nutrition 121, 249–269.

3. Finch NC, Syme HM, Elliott J (2016) Risk Factors for Development of Chronic Kidney Disease in Cats. Journal of Veterinary Internal Medicine 30, 602–610.

4. Chang AR, Anderson C (2017) Dietary Phosphorus Intake and the Kidney. Annual Review of Nutrition 37, 321–346.

5. Dobenecker B, Webel A, Reese S et al. (2017) Effect of a high phosphorus diet on indicators of renal health in cats. J Feline Med Surg 20, 339–343.

6. Pastoor FJ, Van ‘t Klooster AT, Mathot JN et al. (1995) Increasing phosphorus intake reduces urinary concentrations of magnesium and calcium in adult ovariectomized cats fed purified diets. J Nutr 125, 1334–1341.

7. Coltherd JC, Staunton R, Colyer A et al. (2019) Not all forms of dietary phosphorus are equal: an evaluation of postprandial phosphorus concentrations in the plasma of the cat. Br J Nutr 121, 270–284.

8. Dobenecker B, Kienzle E, Schaschl C et al. (2018) Phosphorus source affects renal phosphorus excretion after excessive intake of phosphorus in adult cats. European Society of Veterinary & Comparative Nutrition Congress.

9. Dobenecker B, Hertel-Böhnke P, Webel A et al. (2018) Renal phosphorus excretion in adult healthy cats after the intake of high phosphorus diets with either calcium monophosphate or sodium monophosphate. Journal of Animal Physiology and Animal Nutrition 102, 1759–1765.

10. Finco DR, Barsanti JA, Brown SA (1989) Influence of dietary source of phosphorus on fecal and urinary excretion of phosphorus and other minerals by male cats. Am J Vet Res 50, 263–266.

11. Mack JK, Alexander LG, Morris PJ et al. (2015) Demonstration of uniformity of calcium absorption in adult dogs and cats. J Anim Physiol Anim Nutr (Berl) 99, 801–809.

12. FEDIAF (2019) Nutritional Guidelines For Complete and Complementary Pet Food for Cats and Dogs. March 2019. FEDIAF, The European Pet Food Industry.

13. German AJ, Holden SL, Moxham GL et al. (2006) A Simple, Reliable Tool for Owners to Assess the Body Condition of Their Dog or Cat. The Journal of Nutrition 136, 2031S–2033S.

14. Rokey GJ (1994) Petfood and fishfood extrusion. In The Technology of Extrusion Cooking, pp. 144–189 [ND Frame, editor]. Boston, MA: Springer US.

15. Laflamme DP (2001) Determining metabolizable energy content in commercial pet foods. Journal of Animal Physiology and Animal Nutrition 85, 222–230.

16. Robertson WG, Jones JS, Heaton MA et al. (2002) Predicting the Crystallization Potential of Urine from Cats and Dogs with Respect to Calcium Oxalate and Magnesium Ammonium Phosphate (Struvite). The Journal of Nutrition 132, 1637S–1641S.

17. DeLaurier A, Jackson B, Pfeiffer D et al. (2004) A comparison of methods for measuring serum and urinary markers of bone metabolism in cats. Res Vet Sci 77, 29–39.

18. Geddes RF, Elliott J, Syme HM (2013) The Effect of Feeding a Renal Diet on Plasma Fibroblast Growth Factor 23 Concentrations in Cats with Stable Azotemic Chronic Kidney Disease. Journal of Veterinary Internal Medicine 27, 1354–1361.

19. Williams T, Elliott J, Syme H (2012) Calcium and phosphate homeostasis in hyperthyroid cats–associations with development of azotaemia and survival time. Journal of Small Animal Practice 53, 561–571.

20. Finch NC, Syme HM, Elliott J et al. (2011) Glomerular filtration rate estimation by use of a correction formula for slope-intercept plasma iohexol clearance in cats. American Journal of Veterinary Research 72, 1652–1659.

21. R Development Core Team (2017) R: A language and environment for statistical computing.

22. Bates D, Mächler M, Bolker B et al. (2015) Fitting Linear Mixed-Effects Models Using lme4. 2015 67, 48.

23. Hothorn T, Bretz F, Westfall P (2008) Simultaneous Inference in General Parametric Models. Biometrical Journal 50, 346–363.

24. Lew SW, Bosch JP (1991) Effect of diet on creatinine clearance and excretion in young and elderly healthy subjects and in patients with renal disease. Journal of the American Society of Nephrology 2, 856–865.

25. Reynolds BS, Chetboul V, Nguyen P et al. (2013) Effects of Dietary Salt Intake on Renal Function: A 2-Year Study in Healthy Aged Cats. Journal of Veterinary Internal Medicine 27, 507–515.

26. Yamka RM, Friesen KG, Lowry SR et al. (2006) Measurement of arthritic and bone serum metabolites in arthritic, non-arthritic, and geriatric cats fed wellness foods. International Journal of Applied Research in Veterinary Medicine 4, 265.

27. Baker D, Czarnecki-Maulden G (1991) Comparative nutrition of cats and dogs. Annual review of nutrition 11, 239–263.

28. Hendriks BWH, Wamberg S, Tarttelin MF (1999) A metabolism cage for quantitative urine collection and accurate measurement of water balance in adult cats (Felis catus). Journal of Animal Physiology and Animal Nutrition 82, 94–105.

29. McLeland SM, Lunn KF, Duncan CG et al. (2014) Relationship among serum creatinine, serum gastrin, calcium-phosphorus product, and uremic gastropathy in cats with chronic kidney disease. Journal of veterinary internal medicine 28, 827–837.

30. Landau D, Krymko H, Shalev H et al. (2007) Transient severe metastatic calcification in acute renal failure. Pediatr Nephrol 22, 607–611.

31. Smith BHE, Stevenson AE, Markwell PJ (1998) Urinary Relative Supersaturations of Calcium Oxalate and Struvite in Cats Are Influenced by Diet. The Journal of Nutrition 128, 2763S–2764S.

32. Buffington CA, Rogers QR, Morris JG (1990) Effect of diet on struvite activity product in feline urine. American journal of veterinary research 51, 2025–2030.

33. Lekcharoensuk C, Osborne CA, Lulich JP et al. (2001) Association between dietary factors and calcium oxalate and magnesium ammonium phosphate urolithiasis in cats. J Am Vet Med Assoc 219, 1228–1237.

34. David V, Dai B, Martin A et al. (2013) Calcium Regulates FGF-23 Expression in Bone. Endocrinology 154, 4469–4482.

35. Shimada T, Hasegawa H, Yamazaki Y et al. (2004) FGF-23 Is a Potent Regulator of Vitamin D Metabolism and Phosphate Homeostasis. Journal of Bone and Mineral Research 19, 429–435.

